# NEDD8 activity is important for direct antigen MHC class I antigen presentation

**DOI:** 10.1101/2022.02.25.482016

**Authors:** Kartikeya Vijayasimha, Amy L. Leestemaker-Palmer, James S. Gibbs, Jonathan W. Yewdell, Brian P. Dolan

## Abstract

Successful direct MHC class I antigen presentation is dependent on the protein degradation machinery of the cell to generate antigenic peptides which can be loaded onto MHC class I molecules for surveillance by CD8^+^ T cells of the immune system. Most often this process involves the ubiquitin-proteasome system, however other ubiquitin-like (UBL) proteins have also been implicated in protein degradation and direct antigen presentation. Here, we examine the role of neuronal precursor cell-expressed developmentally down-regulated protein 8 (NEDD8) in direct antigen presentation. NEDD8 is the UBL with highest similarity to ubiquitin and fusion of NEDD8 to the amino-terminus of a target protein can lead to the target proteins degradation. We find that appending NEDD8 to the N-terminus of the model antigen ovalbumin resulted in degradation by both the proteasome and autophagy protein degradation pathways, but only proteasomal degradation, involving the proteasomal subunit NEDD8 ultimate buster 1 (NUB1), resulted in peptide presentation. When directly compare to ubiquitin, NEDD8-fusion was less efficient at generating peptides. However, inactivation of the NEDD8-conugation machinery by treating cells with MLN4924, inhibited the presentation of peptides from Defective Ribosomal Products (DRiPs) derived from a model antigen. These results demonstrate that NEDD8 activity in the cell is important for direct antigen presentation, but not by directly targeting proteins for degradation.

## Introduction

MHC class I molecules are present on the surface of all nucleated cells and are involved in alerting cytotoxic T cells to intracellular infections and oncogenic transformation by presenting an array of endogenous peptides on the cell surface (1, 2). Proteins synthesized in the cytoplasm of the antigen presenting cell undergo proteasomal degradation giving rise to peptides that are loaded onto MHC class I molecules in the endoplasmic reticulum (3). Peptide loading stabilizes the MHC class I molecule that is then transported to the cell surface via the trans-Golgi network (4). Circulating CD8+ T cells scan the MHC class I-peptide repertoire for signs of cognate peptide antigens for which they are specific. Peptide generation is a continuous process and the MHC class I-peptide repertoire provides a constantly updated “view” of the proteomic landscape of the cell (5).

The peptide repertoire generated inside a cell consist of peptides from normal protein turnover within the cell as well as newly synthesized peptides (6). The defective ribosomal product (DRiP) hypothesis of peptide generation identifies post-translationally unstable peptides as being responsible for the generation of the majority of MHC class I peptides (7). A growing body of evidence identifies numerous mechanisms of DRiP generation including post-translationally misfolded or mis-targeted proteins, mistranslated or cryptic translation products, non-canonical translation, and dysregulated protein production resulting disproportional accumulation of polypeptides in the cytoplasm (8). A major merit of the DRiP hypothesis is its ability to account for peptide generation during normal cell function as well as during dysregulation of protein homeostasis.

The ubiquitination of substrates is an integral step for the degradation most proteins that results in the generation of peptides for MHC class I presentation. The addition of multiple Lysine-48 linked ubiquitin molecules to a substrate, is a potent sign for proteasomal degradation, and is the most well characterized degron for proteasomal degradation (9). Mono-ubiquitination (10), especially at the N-terminus (11–13) of substrates has also been shown to be capable of causing proteasomal degradation. Several groups have utilized this cellular property to generate ubiquitin-fusion degradation (UFD) constructs to generate rapidly degraded antigens which leads to increased levels of peptide-MHC complexes and better CD8 T cell responses (14–21). In addition to ubiquitin, a number of ubiquitin like proteins (UBLs) are also capable of causing the proteasomal degradation of substrates. The N-terminal fusion of the UBL FAT 10 (22) to substrates caused the rapid degradation of substrates at the proteasome and resulted in the presentation of peptides derived from conjugates (23). However the fusion of another UBL, interferon stimulated gene 15 (ISG15), to a viral substrate did not result accelerate degradation of substrates, though the presentation of specific epitopes derived from the viral substrate was markedly enhanced (24).

Our previous work with NEDD8 revealed that the fusion of NEDD8 to GFP caused the rapid degradation of GFP (25). This phenomenon was dependent on the proteasomal shuttling factor NEDD8 ultimate buster 1 (NUB1). Further, we showed that a large proportion of NEDD8 modified GFP was degraded lysosomally via the autophagic pathway. The fusion of NEDD8 to a substrate caused a remarkable reduction of the half-life of the substrate. Taken together, these data indicate that NEDD8 likely plays an important role in the degradation and clearance of substrates.

Here, we further characterize NEDD8’s role as a degron and investigated if NEDD8 mediated degradation resulted in peptides for MHC class I presentation. We created fusion proteins in which NEDD8 was fused with a cytoplasmic form of chicken Ovalbumin (OVA) (26) and compared them with similarly constructed ubiquitin-OVA fusion products. Our results show that while non-cleavable (NC) NEDD8-OVA fusion induced both autophagosomal and proteasomal degradation, only proteasomal degradation resulted in increased SIINFEKL presentation. Further, we showed that while proteasomal degradation of NC NEDD8-OVA was partially dependent on the action of NUB1, NC ubiquitin-OVA underwent polyubiquitination, and subsequently proteasomal degradation. When directly compared to a ubiquitin-fusion, NEDD8 fusion resulted in a less robust presentation of the SIINFEKL peptide. However, we did find that inhibition of NEDD8 activation by treatment with MLN4924 could specifically inhibit the presentation of peptides derived from the DRiP form of a separate model antigen. Taken together, these data indicate an important role for NEDD8 in the maintenance of protein homeostasis and peptide generation for presentation with MHC class I.

## Methods

### Plasmids, Antibodies, and Reagents

All constructs were cloned into the eukaryotic expression vector pCAGGS-IRES-Thy1.1, as previously described (25). All primers were obtained from Integrated DNA Technologies. Cloning was confirmed by Sanger DNA sequencing. Fusion proteins were generated as before (25). DNA sequences encoding fused constructs were produced as Gene blocks (from IDT) containing 5’-EcoR1 sites and 3’-Nhel sites for cloning into pCAGGS-IRES-Thy1.1. All constructs contained a truncated, cytoplasmic form of chicken ovalbumin (OVA) (26), either alone or fused with non-cleavable (NC) forms of NEDD8, or ubiquitin, as shown in Figure 1A. NC NEDD8-OVA was constructed by substituting three of the four terminal glycine residues of NEDD8 to alanine before fusing it in frame with OVA. Similarly, NC Ub-OVA was constructed by changing one of the two terminal glycines of ubiquitin to alanine. After cloning, plasmids were prepared for transfection using the Qiagen HiSpeed Midi Plasmid Purification kit according to the manufacturer’s instructions. The following mouse monoclonal antibodies were utilized: anti-Thy1.1 (clone HIS51, eBiosciences), anti-p97 (clone 58.13.3, Fitzgerald), anti poly-ubiquitin (clone FK2, Enzo Life Sciences). The following rabbit polyclonal antibodies were also used: anti-OVA (Sigma, C6534), anti beta-actin (Bethyl Laboratories, A300-485A, Lot# 3), anti-NUB1 (Cell Signaling, 14810S, Lot#1), and anti-NEDD8 (Cell Signaling 2745S, Lot#3). Infrared dye-coupled secondary antibodies (goat anti-mouse or goat anti-rabbit) were from LI-COR. The ubiquitin E1 inhibitor MLN7243 (ChemieTek), and NEDD8 E1 inhibitor MLN4924 (Millipore) were dissolved in DMSO and used at a final concentration of 5 μM. MG132 (Calbiochem) was dissolved in DMSO and used at 10 μm for proteasomal inhibition of cells. 3-methyladenine (3MA) was from EMD Millipore, dissolved in DMSO and used at a final concentration of 50 μM. Bafilomycin (Baf) (Tocris) was dissolved in DMSO and used at a final concentration of 0.1uM. Shield-1 was obtained from Clontech and was used at a final concentration of 2.5 μM.

**Figure 1.**
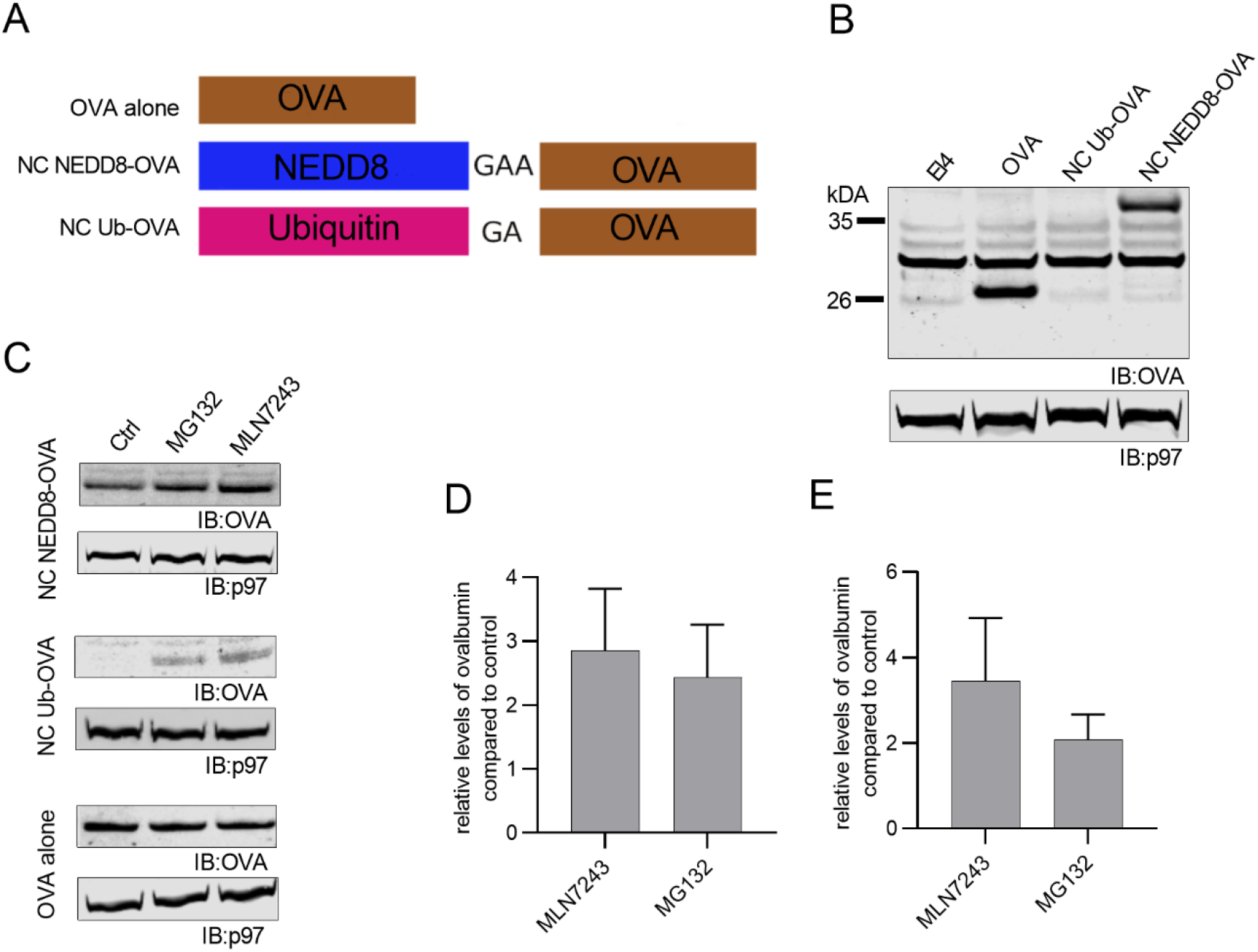
NC NEDD8-OVA and ubiquitin-OVA constructs are degraded in a proteasome- and ubiquitin-dependent manner. *A*. Schematic representation of the constructs including NEDD8 and ubiquitin proteins where the normal C-terminal glycine residues are mutated to alanine and fused with a truncated, cytoplasmic form of ovalbumin (OVA). Both fusion proteins and truncated OVA were cloned into the pCAGGS IRES Thy1.1 expression plasmid and used to stably transfect EL4 cells. *B*. Western blot of EL4, EL4/ova, EL4/NC Ub-OVA and EL4/NC NEDD8-OVA cell lysates. Following SDS-PAGE and blotting onto nitrocellulose, membranes were probed with either rabbit poly-clonal antibodies against OVA or rabbit poly-clonal antibodies against actin and appropriate secondary antibodies. *C*. Western blot of EL4/NC NEDD8-OVA, EL4/NC Ub-OVA and EL4/ OVA cells were treated with either DMSO (mock), MG132, or MLN7243 prior to lysis. *D*. Western blot densitometry was used to determine the relative increase in the OVA signal after MG132, or MLN7243 treatment and is plotted as ratio of OVA in treated samples compared to controls for NC NEDD8-OVA and for NC Ub-OVA. Values represent the mean of multiple independent experiments and error bars represent standard error of the mean.

### Cell Culture, Transfection, and siRNA depletion

L/K^b^, EL4, EL4/NC NEDD8-ova, EL4/ova and EL4/SCRAP-GFP cell lines have been previously described (5, 25, 27). All EL4-derived cells were grown in RPMI 1640 supplemented with 10 mM HEPES, Glutamax, and 7.5% fetal calf serum (all from Thermo Fisher) and were incubated at 37°C and 6% CO_2_. L/K^b^ was cultured in DMEM supplemented with 7.5% FCS and Glutamax. The creation of EL4 cell lines expressing NC Ub-OVA recombinant proteins was accomplished by the transfection of the pCAGGS-IRES-Thy1.1 NC Ub-OVA plasmid followed by magnetic sorting. Briefly, 1 × 10^6^ cells were transfected with 8 μg ScaI-linearized plasmid using the Amaxa Nucleofector (Lonza) SF kit on program DS-113. Transfected cells were allowed to recover for two days and were then labelled at 4°C with anti-Thy1.1 antibody (clone HIS51, eBiosciences) for 30 minutes in PBS buffer containing 0.1% BSA (Amresco). Cells were then washed in buffer and incubated with anti-mouse IgG microbeads beads (Miltenyi Biotech) for 30 min at 4°C, washed, and resuspended in 1 ml buffer. Thy1.1 positive cells were selected by passing cells over LD columns using the MidiMACS separator system per the manufacturer’s instructions. Eluted cells were added to warm media containing antibiotics. Sorted cells were cultured for 48 −72 hours before being sorted again. This process is repeated until a population with > 90% Thy1.1 expression was obtained. For siRNA mRNA interference, cells were harvested by centrifugation and transfected with 1 μM mouse Nub1 (Dharmacon, J-050871-09), or scrambled siRNA (as a negative control) using the Amaxa Nucleofector as described above. Transfected cells were incubated in growth media for 48 hours before being processed for Western blot analysis.

### Flow cytometry

For flow cytometry, cells were collected by centrifugation and washed with Hank’s balanced salt solution (HBSS) containing 0.1% BSA, before being resuspended in 0.1% BSA/HBSS containing isotype control, or PE-Cy5.5 coupled anti-Thy1.1 antibodies (clone HIS51, eBiosciences), or APC coupled 25-D1.16 antibodies (Invitrogen) and incubated for 30 minutes at 4°C. After incubation, cells were washed and resuspended in 0.1% BSA/HBSS before being analyzed with an Accuri C6 benchtop flow cytometer (BD Biosciences). Appropriate filter sets were used to detect APC fluorescence or PE-Cy5.5 fluorescence. All analysis, including determining the mean fluorescence intensity (MFI) for the sample, was conducted using BD Accuri software.

### Antigen Presentation Assays in stable cell lines

Direct MHC class I antigen presentation in stably transfected EL4 cells bearing the SIINFEKL peptide from chicken ovalbumin were conducted by first acid washing cells in ice-cold citric acid buffer (pH3.0) at ~ 2 × 10^7^ cell/ml for 2 minutes. Cells were washed in excess media and returned to culture with the appropriate chemical inhibitors for the indicated times. At the completion of the incubation period, cells were harvested and stained with APC-coupled 25-D1.16 at 0.2 μg/ml for 30 minutes at 4° C and analysed by flow cytometry. Staining was done in quadruplicate and the MFI for each treatment was calculated by averaging the MFI of each sample analyzed. The Shield-controlled Recombinant Antigenic Protein (SCRAP) system is a powerful tool for measuring peptide presentation from three different forms of the same precursor protein: DRiPs, retirees, and normal protein turnover (or non-DRiP). The method for quantifying peptide presentation from this system has been used extensively and previously published (27–32). Briefly, EL4/SCRAP-mCherry cells are washed in citric acid buffer and then re-cultured in either ethanol (negative control), Shield-1, or MG132 for six hours, stained with APC-coupled 25D-1.16 mAbs and analysed by flow cytometry. Quantification of DRiP presentation is achieved by subtracting the MFI of MG132 treated cells from the MFI of Shield-1 treated cells. Quantification of normal protein, or non-DRiP presentation, is made by subtracting the MFI of Shield-1 treated cells from ethanol treated cells. To measure retiree presentation, EL4/SCRAP-mCherry cells are first treated with either Shield-1 or ethanol overnight. The next day the cells are washed, then acid-stripped, then returned to culture and cultured for two hours. An aliquot of cells is stained with 25D-1.16 prior to re-culture after acid washing to determine the level of K^b^-SIINFEKL complexes at time 0. Following two hours of recovery, cells are stained with 25D-1.16 mAbs and analyzed by flow cytometry. The peptide contribution from retirees is determined by subtracting the staining at time 0 for both ethanol and Shield-1 treated cells and then subtracting the resultant ethanol treated MFI values from Shield-1 treated values. In some experiments MLN4924 or MG132 is added to experiments immediately after acid washing.

### Vaccinia virus generation, infection, and antigen presentation

Recombinant vaccina viruses (rVV) were generated as previously described (33). Briefly, the ova-containing gene block fragments were PCR amplified with primers to add a SalI/NotI sites and PCR prodcuts were subsequently cloned in pSC11. CV1 cells were infected with wildtype vaccinia virus and then transfected with pSC11-ova constructs. Recombinant viruses were selected by infecting on TK-cells and plaques identified by beta-galactosidase activity. Three rounds of screening were done to select rVV and stocks created by infecting TK-cells and harvesting rVV from lysed cells. For antigen presentation assays, L/K^b^ cells were washed in HBSS/BSA and resuspended to 2 × 10^7^ cells/ml in HBSS/BSA and infected with rVV at an MOI for 10 for 30 minutes at 37° C with occasional agitating. Cells were washed in media and returned to culture at 10^6^ cells/ml. Aliquots were harvested at indicated times at kept at 4° C until the completion of the experiment when all cells were stained with 25-D1.16 and analyzed by flow cytometry.

### Western Blotting

One million cells were collected by centrifugation and resuspended in cold lysis buffer (1% Triton X-100 in PBS) containing protease inhibitors (Complete protease inhibitor cocktail, Roche) and was incubated on ice for 30 minutes. The solution was centrifuged at 11,000 x g (at 4°C) for 5 minutes, and supernatant removed to a fresh tube containing Bolt LDS buffer (4X) with 10 μM DTT. The cell lysate was heated to 95°C for 10 minutes and was resolved by SDS-PAGE using a 4-12% Bolt Bis-Tris gel (Thermo Fisher) and transferred to a nitrocellulose membrane (Thermo Fisher). Membranes were then transferred to a container with a 5% solution of dehydrated milk in TBST (Tris-buffered saline, 0.1% Tween 20) and was placed on a rocker for 40 minutes for blocking. Membranes were then washed with TBST and incubated in Primary antibody solution (1:1000-1:5000 in TBST containing 0.5% milk) overnight. Secondary antibody solution (1:10,000 in TBST containing 0.5% milk) was added to the membrane after washing and membranes incubated for an hour, followed by two five-minute washes in TBST and one rinse in deionized water. Membranes were analyzed using an infrared imager (LI-COR Odyssey) and blots quantified by densitometry using instrument software (Image Studio Lite, LI-COR Biosciences). Relevant bands were determined based upon either predicted size of the protein or by its presence/absence in control experiments. All western blot images were cropped to show relevant bands. Adjustments to the brightness or the contrast were applied uniformly to the entire image. Such alterations did not change the intensity values obtained by the instrument software.

### TUBE-isolation Assay

The polyubiquitinated status of proteins in cells was determined using the Tandem Ubiquitin Binding Entities kit (TUBE, Life Sensors). Detection of polyubiquitination was carried out as per the manufacturer’s instructions. Briefly, 3.5 × 10^7^ cells were cultured for 2 hours in the presence of MG132 (10 μm) and then washed in cold PBS. Cells were spun down again and the pellet resuspended in 1.25 ml cold lysis buffer (PBS containing 1% TX-100, 1 mM NEM, and Complete Protease Inhibitor from Roche). Cells were lysed for 30 minutes on ice. The lysate was then centrifuged at 10,000 x g for 10 minutes at 4°C and a portion of supernatant saved as the pre-clear analysis. The TUBE beads were washed twice and resuspended in cold lysis buffer. For the isolation of poly-ubiquitinated proteins, 125 μl of prepared TUBEs were added to the cell lysate, and incubated at 4°C for 1 hour while being constantly mixed. TUBE beads were collected by centrifugation at 5,000 x g and an aliquot of supernatant was saved for post-clear analysis and the rest discarded. The beads were washed three times with 700 μl cold lysis buffer and resuspended in 40 μl lysis buffer, 20 μl BOLT sample buffer, and 2 μl of 1M DTT and heated at 95 °C for 10 minutes. The solution was centrifuged and the supernatant analyzed by western blot analysis.

### Statistical Analysis

All statistical analysis was conducted using Prism (Graphpad). Changes in levels of a given protein from controls were determined by calculating the ratio of protein in treated samples to controls. The mean value from at least three independent experiments was analysed for statistical significance using a one-sample t-test assuming a theoretical mean value of 1 (no fold-change in protein). Unpaired t-tests were used to compare differences in mean values in fluorescence intensities for K^b^-SIINFEKL levels from at least three technical replicates when comparing different treatments. Changes were considered statistically different at a P value <0.05.

## Results

### Non-cleavable NEDD8 and Ubiquitin ovalbumin constructs are degraded in a proteasome- and ubiquitin-dependent manner

In order to study the effect of non-canonical NEDD8 binding on antigen presentation we created three OVA fusion proteins (Figure 1A), similar to previously described GFP constructs (25). OVA consists of a truncated, cytoplasmic form of chicken ovalbumin encoded by amino acids 146-386 (26). Non-cleavable (NC) forms of NEDD8-OVA and ubiquitin-OVA, henceforward referred to as NC NEDD8-OVA and NC Ub-OVA respectively, were generated by fusing the open reading from of OVA to the c-terminal end of either NEDD8 or ubiquitin and mutating the di-glycine (ubiquitin) or 4-glycine repeat (NEDD8) to contain alanine substitutions. All constructs were cloned into the eukaryotic expression vector pCAGGS IRES Thy1.1. Stable cell lines were generated by transfecting EL4 cells with these constructs and sorting for Thy1.1 expression using magnetic beads. Multiple rounds of sorting were carried out until equivalent levels of Thy1.1 expression (>90%) was seen in all cell lines via flow cytometry. We then examined each cell line for the presence of ovalbumin by western blot. Several non-specific, antibody reactive bands, were detected in EL4 cells alone (Figure 1B), however, a discernible band of ~ 26kDa was detected in EL4/OVA cell lysates, which corresponds to the predicted size of the cytosolic form of ovalbumin (Figure 1B). A less intense band of ~ 34kDa was detected in the lysates of EL4/NC NEDD8-OVA cell lysates which corresponds to the size of cytosolic ovalbumin fused with NEDD8. No OVA-specific bands were readily observable in EL4/NC Ub-OVA cell lysates (Figure 1B). The lack of protein in the EL4/NC Ub-OVA cells may result from the rapid degradation of the protein due to the ubiquitin fusion. To test this hypothesis, we treated EL4/NC NEDD8-OVA, EL4/NC Ub-OVA cells and EL4/OVA with either MG132 to prevent proteasome degradation or MLN7243, a potent inhibitor of the ubiquitin conjugation system. Following treatment, cells were cultured for two hours prior to lysis and western blot analysis. Inhibiting either the proteasome or the ubiquitin conjugation system, resulted in increased levels of both NC NEDD8-OVA and NC Ub-OVA proteins in cell lysates (Figures 1C). Levels of cytosolic ovalbumin in EL4/OVA cells remained relatively unchanged. The calculated ratio of OVA levels in EL4/NC NEDD8-OVA (Figure 1D) and EL4/NC Ub-OVA (Figure 1E) following treatment with either MG132 or MLN7243 to OVA levels in control cells from multiple, independent, experiments is shown and is statistically different from a theoretical mean of 1 for all treatments (P <0.05). Taken together, these data indicate that the fusion of NEDD8 or ubiquitin to cytosolic ovalbumin results in targeting to the proteasome and renders the protein rapidly degraded, consistent with previous reports.

### Ubiquitin fusion increase direct antigen presentation compared to NEDD8 fusion

Data presented in Figure 1 demonstrate that direct conjugation of NEDD8 to a protein can lead to rapid degradation. We hypothesize that the rapid degradation of NEDD8-conjugated ovalbumin would lead to enhanced presentation of peptides from ovalbumin. To test this, we generated recombinant vaccinia viruses containing genes encoding either cytosolic OVA, NC NEDD8-OVA, or NC Ub-OVA. Recombinant viruses were then used to infect L/K^b^ cells and the presentation of the ovalbumin peptide SIINFEKL by the MHC class I molecule K^b^ was monitored using a monoclonal antibody specific for the K^b^-SIINFEKL complex. Use of rVV is desirable for comparing antigen presentation from different substrates for several reasons including synchronizing the time of antigenic protein synthesis, ensuring an equivalent level of substrates synthesized, and utilization of a homogenous population of antigen presenting cells at the time of infection. Fusion of cytosolic ovalbumin to ubiquitin resulted in a rapid increase in antigen presentation, similar to data reported elsewhere (14). However, conjugation of NEDD8 was not nearly as efficacious at generating K^b^-SIINFEKL complexes (Figure 2A) compared to ubiquitin conjugation. Levels of K^b^-SIINFEKL were still greater than what was observed when cells were infected with rVV expressing cytosolic ovalbumin alone, indicating that NEDD8 fusion can partially enhance presentation over what would be considered background presentation. We then compared viral antigen presentation of non-cleavable and cleavable forms of the substrates. Additional rVV were generated which expressed either ubiquitin or NEDD8 fusion proteins, but unlike the non-cleavable forms of the protein, the glycine motifs remained intact, allowing ubiquitin and NEDD8 processing enzymes to remove the cytosolic ovalbumin from either ubiquitin or NEDD8. Comparison of K^b^-SIINFEKL generation from the ubiquitin conjugated ovalbumin constructs demonstrate enhanced peptide presentation from the non-cleavable form of the substrate (Figure 2B), again consistent with previous reports indicating that irreversible fusion of ubiquitin to a substrate enhances peptide presentation. However, presentation of the SIINFEKL peptide was essentially identical when non-cleavable and cleavable forms of NEDD8 conjugated ovalbumin were examined (Figure 2C). These data indicate that unlike ubiquitin-conjugation, modification of a substrate with NEDD8 does not lead to robust peptide-presentation.

**Figure 2.**
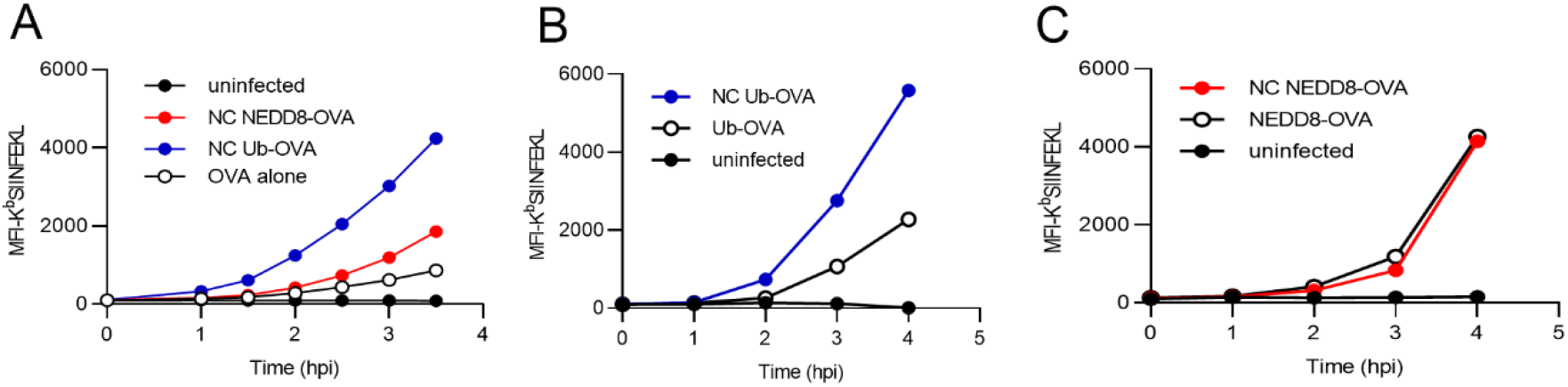
Increased efficiency of peptide presentation from ubiquitin-fused OVA compared to NEDD8-fused OVA. Recombinant vaccinia viruses containing genes encoding cytosolic ovalbumin and either non-cleavable, or cleavable NEDD8 or ubiquitin conjugated forms of cytosolic ovalbumin were generated and used to infect L/K^b^ cells. K^b^-SIINFEKL presentation on infected cells was gauged at the indicated times by using 25-D1.16 mAbs. *A*. Comparison of mean fluorescence intensities of K^b^-SIINFEKL after infection in L/K^b^ cells following infection with rVV expressing NC Ub-OVA (blue, solid circle), NC NEDD8-OVA (red, solid circle), OVA alone (black, empty circle), and uninfected cells (black, solid circle). *B*. Comparison of K^b^-SIINFEKL presentation in NC Ub-OVA (blue, solid circle) and Ub-OVA (black, empty circle). *C*. Comparison of K^b^-SIINFEKL presentation from NC NEDD8-OVA (red, solid circle), and NEDD8-OVA (black, empty circle).

### Autophagy inhibition affects NC NEDD8-OVA degradation but does not result in increased presentation

Our previous work indicated that modification of GFP with non-cleavable NEDD8 resulted in the lysosomal degradation of a substantial portion of NC NEDD8-GFP, via the autophagy pathway. In order to check if the non-cleavable attachment of NEDD8 or ubiquitin to OVA resulted in autophagy-mediated lysosomal degradation, we treated EL4/NC NEDD8-OVA, EL4/NC Ub-OVA, and EL4/OVA alone cells with the autophagy inhibitors, 3-methyladenine (3MA), at a concentration of 50mM for 5 hours, and Bafilomycin, at a concentration of 0.1 μM for 6 hours. Degradation of OVA after autophagy inhibition was measured by western blotting. As previously observed for NC NEDD8-GFP we detected an increase in NC NEDD8-OVA following treatment with autophagy inhibitors in EL4/NC NEDD8-OVA cells (Figure 3A). However, treatment of EL4/NC Ub-OVA cells with autophagy inhibitors did not increase the levels of the non-cleavable Ub-OVA protein (Figure 3A), in contrast to what was observed when cells were treated with proteasome inhibitors (Figure 1C). There was no change in the level of cytosolic ovalbumin in EL4/OVA alone cells treated with either autophagy inhibitor (Figure 3A). The ratio ovalbumin present in cells following autophagy compared to control cells in NC NEDD8-OVA is plotted in figure 3B plots and is statistically significant (P < 0.01). These data demonstrate that NEDD8 conjugation, but not ubiquitin conjugation, leads to degradation of a subset of proteins by the autophagy pathway.

**Figure 3.**
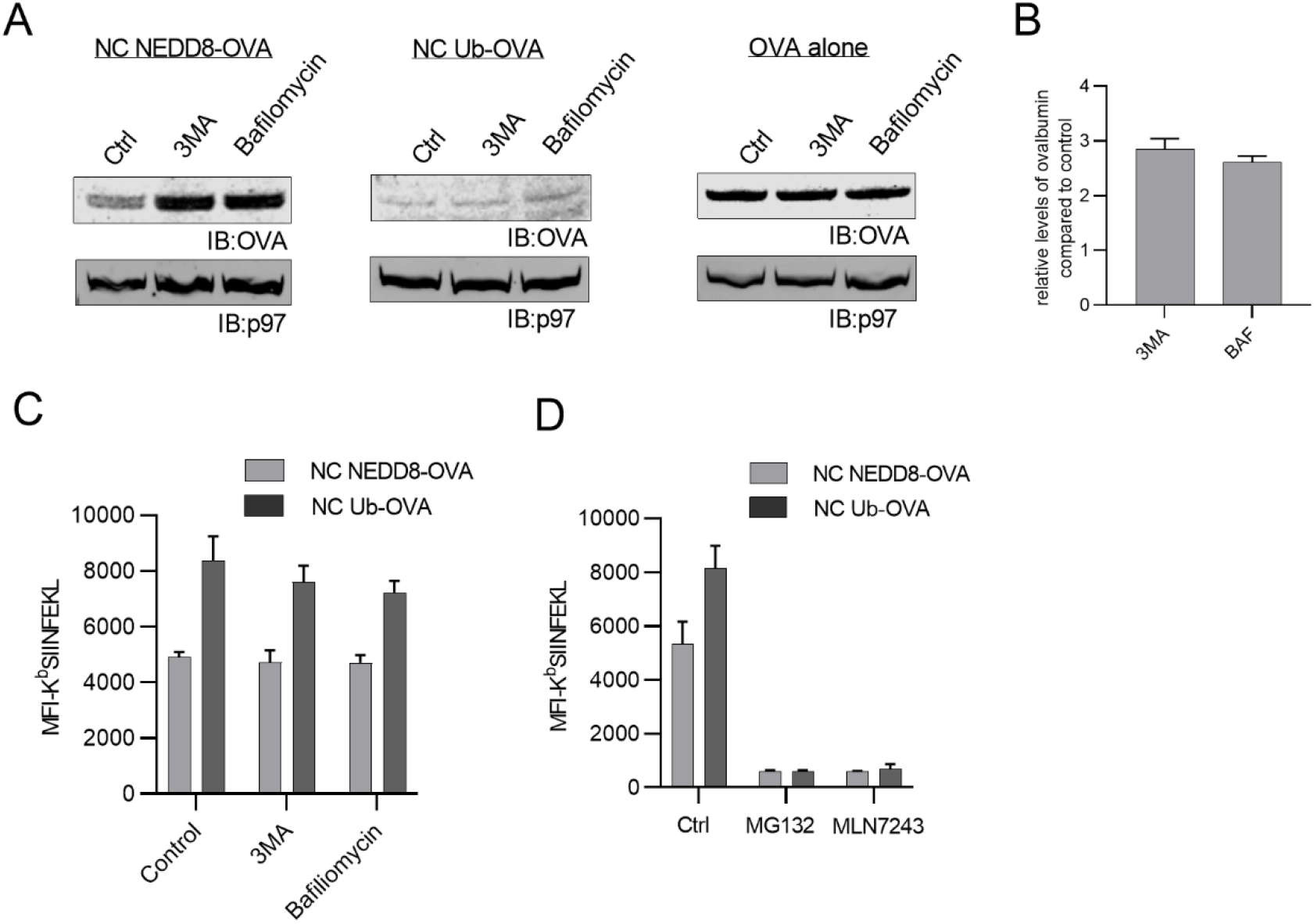
Non-cleavable NEDD8-OVA undergoes autophagosomal degradation. *A*. EL4/NC NEDD8-OVA, EL4/NC Ub-OVA, and EL4/OVA cells treated with the autophagy inhibitors 3-MA or bafilomycin and analyzed by western blot analysis for ovalbumin. *B*. The ratio of ovalbumin in treated samples compared to controls as determined by western blot densitometry in EL4/NC NEDD8-OVA cells after autophagy inhibition, averaged from three independent experiments. *C*. Flow cytometry analyses to measure mean fluorescence intensity of K^b^-SIINFEKL in EL4/NC NEDD8-OVA and EL4/NC Ub-OVA cells after autophagy inhibition. *D*. Flow cytometry analyses to measure mean fluorescence intensity of K^b^-SIINFEKL in EL4/NC NEDD8-OVA, and EL4/NC Ub-OVA after inhibition of the proteasome (MG132) or the ubiquitin-conjugation pathway (MLN7243). All values represent the mean of three independent experiments and error bars represent standard error.

Having established that a portion of non-cleavable NEDD8-OVA was degraded by the autophagy pathway, we sought to determine what effect inhibiting autophagy would have on antigen presentation. EL4/NC NEDD8-OVA cells were washed in a mild citric acid buffer to remove existing peptide-MHC complexes from the cell surface and cultured for 6 hours in the presence of 3-MA, bafilomycin, or DMSO as a control. After staining cells with the K^b^-SIINFEKL-specific monoclonal antibody, cells were analyzed by flow cytometry to determine the levels of peptides presented. Inhibiting autophagy had no effect on the presentation of SIINFEKL peptide from the non-cleavable form of NEDD8-OVA, or the non-cleavable forms of Ub-OVA (Figure 3C). As an additional control, we cultured acid-washed cells in either MG132 or MLN7243 for two hours and measured peptide-presentation. Inhibiting either proteasome degradation or ubiquitin conjugation almost completely abolished antigen presentation, not only in EL4/NC NEDD8-OVA cells but also in EL4/NC Ub-OVA cells (Figure 3D). It is also worth noting that while we cannot directly compare the efficacy of presentation between the different cell lines, the pattern we observed in rVV infection was maintained in stably transfected EL4 cells, namely increased peptide presentation from NC Ub-OVA compared to NC NEDD8-OVA. These data indicate that the attachment of NEDD8 to OVA resulted in a significant proportion of NC NEDD8-OVA being degraded via autophagy, while the attachment of ubiquitin did not lead to autophagy degradation of OVA. Further, these data also indicate that autophagy does not contribute significantly to the presentation of the SIINFEKL peptide from the degradation of NC NEDD8-OVA.

### NC ubiquitin-OVA is polyubiquitinated

The tagging of a substrate protein with multiple ubiquitin moieties is widely regarded as a degron for proteasomal degradation. As the SIINFEKL presentation levels of NC NEDD8-OVA were significantly lower than that of NC Ub-OVA we wanted to check if the polyubiquitination status of NC NEDD8-OVA differed from that of NC Ub-OVA proteins. To determine poly-ubiquitination status we utilized tandem ubiquitin binding entities (TUBEs) beads to isolate poly-ubiquitinated proteins from cell lysates. Cells were treated with MG132 for 2 hours to allow for the buildup of poly-ubiquitin tagged proteins destined for proteasomal degradation and then lysed and mixed with TUBES to precipitation poly-ubiquitinated proteins. The TUBES assay was able to pull down almost all poly-ubiquitinated proteins from lysates as observable by western blotting and staining for ubiquitin (Figure 4A). Staining the TUBE-isolated protein demonstrated very little beta-actin present, suggesting that the protein isolated by TUBES were largely poly-ubiquitinated with very little contamination of non-ubiquitinated proteins. We then analyzed TUBES-isolated proteins for ovalbumin by western blotting. As shown in Figure 4B, a number of non-specific, antibody-reactive, bands were present in EL4 cell lysates. In EL4/ova cells, the only OVA-specific band present corresponded to free cytosolic ovalbumin alone (26kDA) (Figure 4B) while in the EL4/NC NEDD8-OVA cell lysate a band corresponding to the non-cleavable NEDD8-OVA fusion protein was present (34kDA). However, no unique higher molecular weight bands were identified in either EL4/OVA cell lysates or EL4/NC NEDD8-OVA cell lysates. In EL4/NC Ub-OVA cells a higher molecular weight OVA reactive band was detected which corresponded to the size of Ub-OVA plus an additional ubiquitin molecule (Figure 4B, asterisk). Furthermore, this band was dependent on the presence of functional ubiquitin conjugation machinery in the cell as treatment with MLN7243 greatly weakened the signal associated with this band (Figure 4C). Therefore, NC Ub-OVA can be decorated with at least one additional ubiquitin proteins while NC NEDD8-OVA is not.

**Figure 4.**
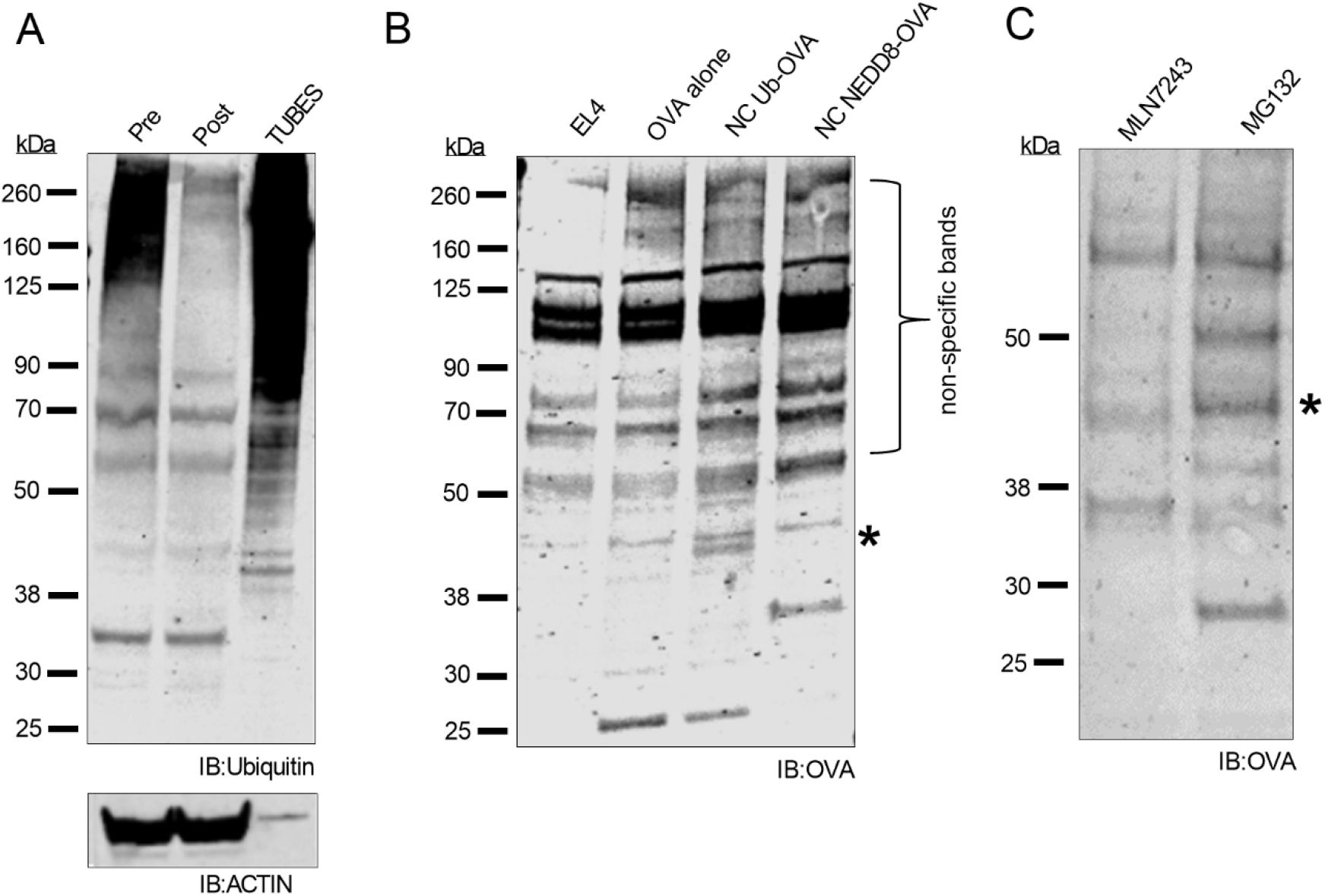
Poly-ubiquitination of non-cleavable Ub-OVA is required for its degradation. Poly-ubiquitnated proteins were isolated from MG132-treated EL4, EL4/ OVA alone, EL4/NC NEDD8-OVA, and EL4/NC Ub-OVA cell lysates using TUBEs and subjected to western blot analysis. *A*. Western Blot analyses of EL4/NC NEDD8-OVA lysates before TUBES treatment (labelled pre-clear), post-pull down (labelled post-clear), and the TUBEs precipitated proteins. Ubiquitinated proteins were identified by staining with the ubiquitin-specific monoclonal antibody FK2. Actin (bottom panel) is shown as a control. *B*. Poly-ubiquitinated proteins were isolated from EL4, EL4/NC NEDD8-OVA, and EL4/NC Ub-OVA cell lysates using TUBES and were subjected to western blot analysis with OVA polyclonal antibodies. Poly-ubiquitinated NC Ub-OVA is marked with an asterisk. *C*. Western blot analysis of EL4/NC Ub-OVA cell lysates after treatment with either MLN7243 or MG132. EL4/NC Ub-OVA cells were treated with either MG132 to allow accumulation of polyubiquitinated proteins, or MLN7243 to prevent ubiquitination of OVA. Lysates were subjected to western blot analysis for OVA. The ~44 kDa band corresponding to polyubiquitinated OVA present in MG132 treated lysates is marked with an asterisk.

### NUB1 is necessary for degradation of NEDD8 conjugated, but not Ubiquitin conjugated proteins

We have previously demonstrated that proteasomal mediated degradation of non-cleavable NEDD8 GFP required NUB1 (25). To test if NUB1 played a role in the degradation of NC NEDD8 OVA, NC Ub-OVA, or OVA alone, we transfected EL4/NC NEDD8-OVA, EL4/NC Ub-OVA, or EL4/OVA cells with either siRNA oligomers targeting NUB1 or scrambled oligomers acting as a negative control. After transfection, cells were processed for western blotting. NUB1 expression was greatly reduced in all cell lines following transfection of NUB1 siRNA nucleotides (Figure 5A top panels). NUB1 silencing resulted in accumulation of NC NEDD8-OVA, while no change was detected by western blotting in levels of NC Ub-OVA or OVA alone (Figure 5A). Quantification of western blot densitometry showed an increase in NC-NEDD8 OVA after NUB1 KD (Figure 5B, P < 0.05). No significant increase was seen in EL4/OVA cells or EL4/NC Ub-OVA cells. Because NUB1 can associate with the proteasome and proteasome inhibition abolishes presentation of peptides from NC NEDD8-OVA, we tested what role NUB1 depletion would have on peptide presentation. NUB1-depleted cells were acid washed to strip existing peptide-MHC class I complexes from the cell surface and cultured in regular media under normal conditions for 4 hours. Cells were then stained with 25-D1.16 antibodies and mean fluorescence intensity of K^b^-SIINFEKL measured by flow cytometry. Presentation of peptides in NUB1 depleted cells was normalized to cells treated with scrambled siRNA controls. EL4/NC NEDD8-OVA cells showed a greater than 50% reduction in SIINFEKL presentation after NUB1 knockdown (Figure 5C, P <0.001), which did not occur in either EL4/OVA cells or EL4/NC Ub-OVA cells. These data indicate that NUB1 does play a role in the degradation and subsequent presentation of SIINFEKL in EL4/NC NEDD8-OVA cells.

**Figure 5.**
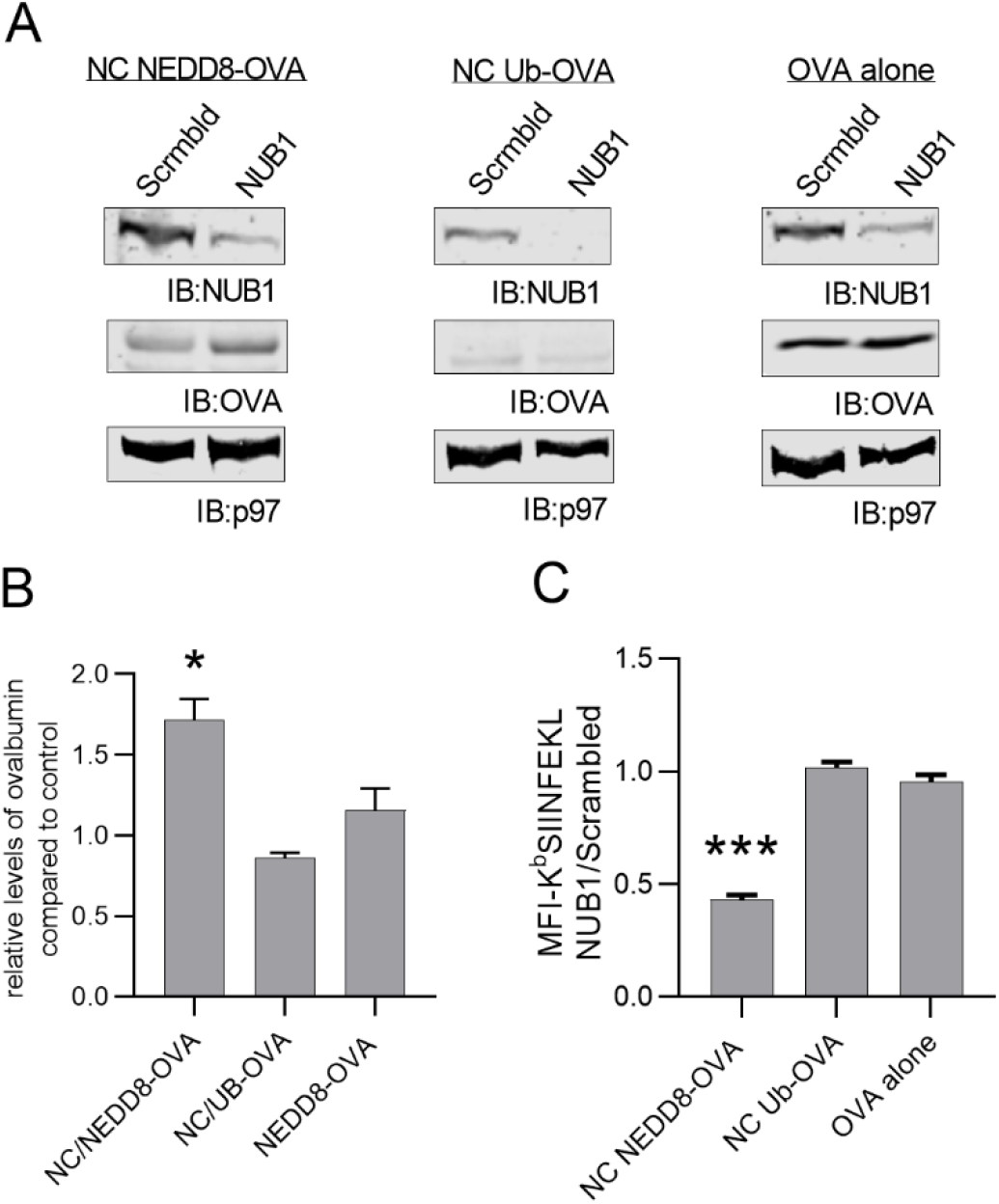
NUB1 is necessary for degradation and direct antigen presentation of NEDD8 conjugated OVA. *A*. EL4/NC NEDD8-OVA, EL4/NC Ub-OVA, or EL4/OVA cells were transfected with either siRNA oligomers targeting NUB1 or scrambled oligomers and processed for western blot analysis for NUB1 (top panels) and OVA (middle panels). All membranes were blotted for p97 (bottom panels) as a loading control. *B*. The ratio of ovalbumin in siRNA depleted cells, as determined by densitometry, is compared to control cells transfected with scrambled siRNA. *C*. Flow cytometric analysis for mean fluorescence intensity of K^b^-SIINFEKL in EL4/NC NEDD8-OVA, EL4/NC Ub-OVA, or EL4/OVA cells after NUB1 silencing. Changes in MFI K^b^-SIINFEKL are shown as a ratio of fluorescence in NUB1 silenced cells over cells transfected with scrambled siRNA. Values represent mean from three independent experiments. The mean values is statistically different from the theoretical value of 1 (*P < 0.05, ***P < 0.001).

### Presentation of Peptides from DRiPs is dependent on NEDD8 activation

While the data presented thus far suggest that NEDD8 conjugation to a substrate can lead to degradation, direct conjugation of NEDD8 is not as robust as ubiquitin conjugation for the presentation of peptides via MHC class I. To determine if NEDD8 activation was important for the presentation of peptides from other non-NEDD8 conjugated substrates, we turned to another model substrate termed Shield-Controlled Recombinant Antigenic Protein or SCRAP. SCRAP proteins consist of a modified FKBP12 protein which serves to destabilize the substrate to which it is attached (34). The destabilization domain is fused in frame with the SIINFEKL peptide followed by a reporter protein, in this case mCherry. As SCRAP-mCherry is degraded, the SIINFEKL peptide is liberated for presentation on K^b^ molecules, which can be measured by flow cytometry. The presence of the small molecule Shield-1, allows the protein to fold, prevents its degradation, and allows for gain of function, *i.e*, fluorescence. The use of Shield-1 to stabilize SCRAP-mCherry allows us to control the source of SIINFEKL peptides for MHC class I antigen presentation and independently measure peptide presentation from previously synthesized SCRAP-mCherry being “retired” by the cell, normal protein turnover based upon the predicted half-life of SCRAP-mCherry, and Defective Ribosomal Products (DRiPs) which is a form of SCRAP-mCherry which is inherently refractory to stabilization by Shield-1 treatment. This system has been used multiple times to identify which cellular components control presentation of peptides from their respective sources (27–32).

To determine if NEDDylation was necessary for the presentation of peptides, we acid washed EL4/SCRAP-mCherry cells to remove existing peptide-MHC class I complexes and cultured cells with or without Shield-1 in combination with MLN4924, a potent inhibitor of the E1 activating enzyme for NEDD8 NAE1, for 5 hours. MLN4294 treatment did not appreciably impact the ability of EL4 cells to synthesize SCRAP-mCherry (Figure 6A) nor did it prevent cells from degrading previously synthesized SCRAP-mCherry (Figure 6B). Levels of K^b^-SIINFEKL were determined by flow cytometry and background levels of antigen presentation were determined by examining cells treated with MG132 after acid wash. Presentation of peptides from normal protein turnover of substrates, as measured in the absence of Shield-1, was unaffected by MLN4924 treatment (Figure 6C), however, presentation of peptides from DRiP substrates was dramatically reduced by treatment with MLN4924 (Figure 6C, P < 0.05). Presentation of peptides from retired SCRAP-mCherry proteins was unaffected by MLN4924 treatment (Figure 6C), while preventing retiree degradation by inhibiting the proteasome did prevent retiree presentation. Combined these data indicate that cellular NEDDylation is important for presentation of peptides from DRiP substrates, but does not generally inhibit MHC class I presentation or the presentation of peptides from non-DRiP forms of SCRAP-mCherry

**Figure 6.**
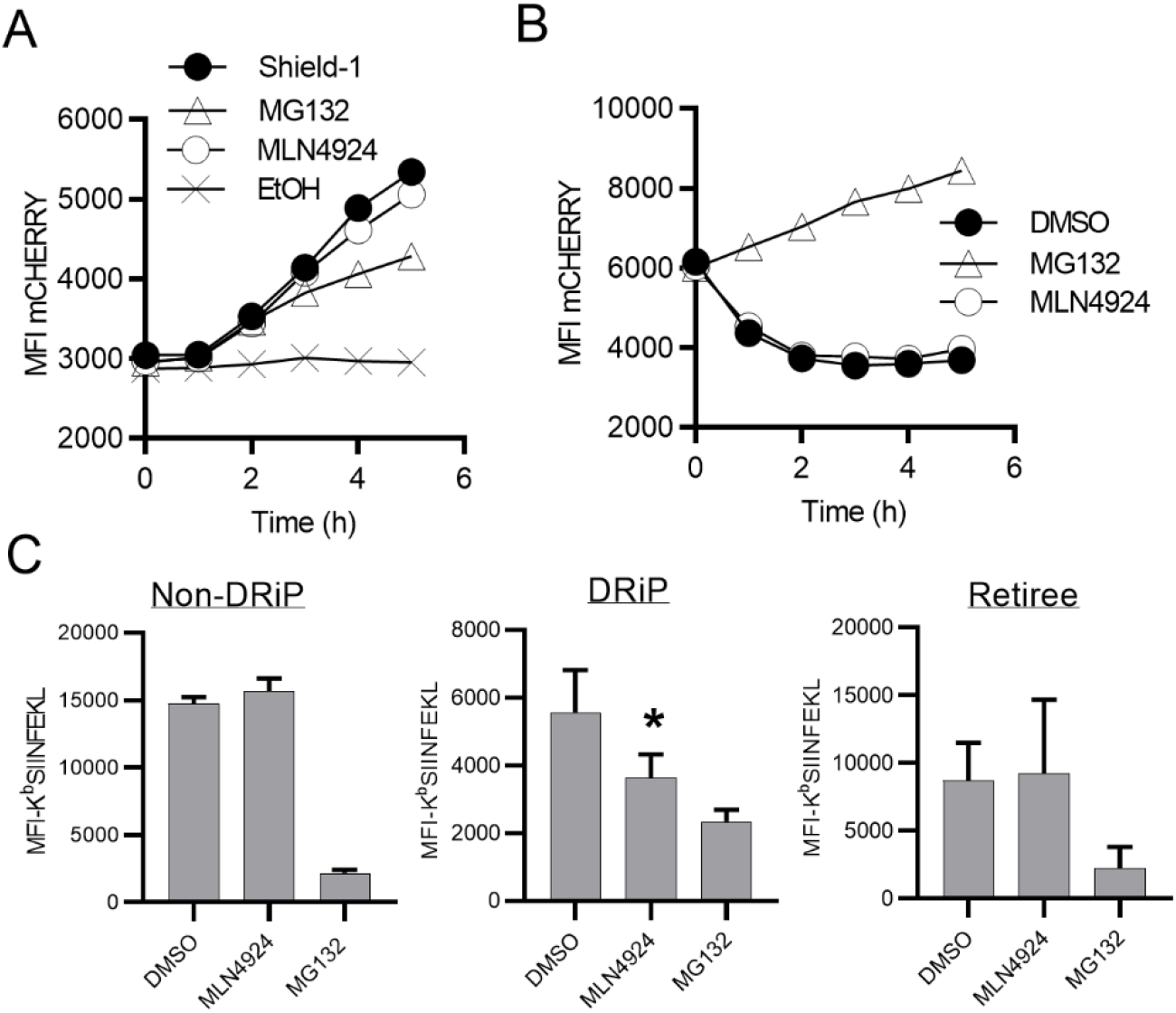
Presentation of peptides from DRiPs is dependent on NEDD8 activation. *A*. EL4/SCRAP mCherry cells were washed in a mild acid buffer to remove existing K^b^-SIINFEKL complexes and cultured in the presence of Shield-1 with DMSO, MLN4924, or MG132 and mCherry fluorescence determined by flow cytometry at indicated times. *B*. EL4/SCRAP mCherry cells were treated with Shield-1 overnight to build up a pool of SCRAP-mCherry, then washed as in *(A)* and cultured without Shield-1 in the presence of DMSO, MLN4924, or MG132. Fluorescence of mCherry was measured at indicated times. *C*. To determine NEDD8’s involvement in peptide presentation from SCRAP, EL4/SCRAP-mCherry cells were acid washed and cultured with or without Shield-1 and either MLN4924 or MG132. Presentation of peptides from Non-DRiP, DRiP, and retiree forms of SCRAP with or without MLN4924 treatment was measured as described in the Materials and Methods section. Only presentation of DRiP substrates following MLN4924 treatments was statistically different from DMSO-treated cells (P < 0.05). MG132 treatment is shown as a negative control.

## Discussion

The ubiquitin fusion degradation (UFD) pathway has been used by multiple groups to enhance our understanding of direct MHC class I antigen presentation and to drive the activation of CD8+ T cells targeting multiple types of pathogens and tumors (14–21). The strategy of genetically manipulating ubiquitin fusion proteins has been extended to the study of other ubiquitin-like proteins (UBLs). For instance, the UBL FAT10, can induce the degradation of its fusion partner, resulting in increased presentation of antigenic peptides (22, 23). For other UBLs, the story is more murky. N-terminal fusion of ISG15, a UBL who’s production is upregulated following viral infection and binds to newly synthesized proteins, does not have an apparent impact on the stability of the protein, though an increase in peptide presentation has been demonstrated (24). In other instances, such as N-terminal fusion of SUMO, there is little to no impact either protein stability or antigen presentation (22). We have recently reported the N-terminal fusion of NEDD8 to a protein induces rapid degradation (25) and in this report sought to determine if the induction of protein degradation by NEDD8 fusion would enhance antigen presentation. By comparing cells infected with rVV expressing the different constructs, we were able to directly compare the effect of NEDD8-fusion with ubiquitin fusion and found that while NEDDylation of the cytosolic form of ovalbumin increased the rate of antigen presentation, it was not as efficient as ubiquitin fusion.

There are several plausible reasons why the presentation of peptides from a NEDDylated protein may not be as efficient when compared to a ubiquitinated protein. From a cell biology standpoint, the N-terminal fusion of NEDD8 to a protein can induce degradation by both the proteasome and via the autophagy pathway (25). Inhibiting autophagy resulted in a partial rescue of NEDD8-ova levels within cells (Figure 3A), however, inhibiting autophagy had no impact on the presentation of peptides from the NEDD8-ova fusion protein (Figure 3C). Given the dependence on proteasome degradation and TAP-dependence for direct MHC class I presentation, it is unlikely that proteins degraded by autophagy serve as a source of peptides for direct presentation, though this pathway may be important for MHC class I cross-presentation by dendritic cells (35, 36). It is therefore likely that the fraction of NEDDylated ovalbumin targeted for degradation in the autophagosome is prevented from donating peptides to the MHC class I pathway, whereas the ubiquitin-fused ovalbumin which is degraded exclusively by the proteasome provides more substrates to the proteasome for degradation.

In addition to shunting a portion of substrates to the autophagy pathway, N-terminal NEDDylation does not appear to induce poly-ubiquitination of NEDD8-OVA, similar to what we have previously reported for NEDD8-GFP (25). In contrast, ubiquitin-OVA did have a distinct band isolated following a TUBES-pull down assay which suggested multiple ubiquitin monomers or chains covalently attached to the substrate. The addition of ubiquitin to the substrate may enhance the substrates degradation. Indeed, despite a similar level of expression as determined by a reporter protein from the bi-cistronic vector used to introduce the fusion protein into cells, levels of NC Ub-OVA were not detectable (Figure 1B) while a low level of NC NEDD8-OVA could be measured in cells. This would suggest that the ubiquitin-OVA construct is degraded at a much faster rate that the NEDD8-ova fusion protein. A faster degradation rate, perhaps induced by poly-ubiquitination, could explain the increased levels of peptide presentation observed in the ubiquitin-ova constructs.

In addition to examining the impact of direct NEDDylation of a substrate on peptide presentation, we also examined the effect of global NEDDylation inhibition on the presentation of peptides from a model substrate whose stability can be post translationally controlled. When cells were treated with MLN4924, the presentation of peptides derived from DRiP substrates was greatly reduced. Inhibiting NEDDylation did not impact the presentation of peptides derived from normal protein turnover, nor did it effect the presentation of peptides from retired proteins, indicating that MLN4924 is not a generalized inhibitor of MHC class I antigen presentation. The model substrate from which the peptide was derived, SCRAP, was also unaffected by MLN4924 treatment as both the synthesis of the protein and its degradation were unaltered by drug treatment. Given that MLN4924 treatment only diminishes DRiP presentation, one could hypothesize that the DRiP form of SCRAP needed to be NEDDylated prior degradation and antigen presentation. However, DRiP presentation is thought to be a highly efficient process whereby few precursor substrates are needed to be degraded for the presentation of peptide antigens (27, 37). If NEDDylation does result in a decrease in antigen presentation efficiency as suggested by the data presented here, it would be unlikely that DRiPs synthesized by the cell are directly modified by NEDD8 prior to their degradation and presentation.

Inhibiting global NEDDylation will also prevent the ubiquitin ligation functions of Cullin-Ring ligases (CRLs), the largest family of E3 ubiquitin ligases. DRiP presentation may therefore rely on the activity of CRLs to identify and ubiquitnate DRiPs prior to degradation, while the retiree and non-DRiP forms of SCRAP are degraded and presented independently of CRLs. If true, this would provide further evidence that DRiPs are treated differently by the cell than other forms of a protein destined for degradation. Previously stabilized SCRAP is degraded independently of MLN4924, suggesting that CRL activity is not needed for SCRAP ubiquitination and degradation by the proteasome, yet the DRiP form of SCRAP, whatever that may be, could be recognized and ubiquitinated by one or more CRLs. Future work should investigate this more closely, especially as MLN4924 is now being used in more clinical trials for the treatment of numerous cancers and many have suggested combinatorial therapies utilizing MLN4924 with T cell checkpoint inhibitors (38, 39). If MLN4924 has an effect on the source of peptides presented by tumor cells, then activated tumor-specific T cells may be exposed to a different peptide repertoire impacting the efficacy of T cell therapies. Clearly, this topic will require careful study.

Finally, it is worth noting that another target of post translational NEDDylation are a subset of ribosomal proteins (40). The exact reason for ribosomal NEDDylation are not fully understood, though it may protect ribosomes from degradation during certain cellular stressors (41, 42).

However, we and others have long-speculated on the presence of an “immunoribosome” which may translate DRiPs and lead to efficient degradation and antigen presentation (43, 44). Perhaps NEDDylation of certain ribosomal proteins would permit this to occur, in which case, MLN4924 treatment would block the ribosomal NEDDylation and prevent DRiP synthesis.

## Abbreviations

3MA: 3-methyladenine
DRiP: Defective ribosomal product
NC: non-cleavable
NEDD8: neuronal precursor cell-expressed developmentally down-regulated protein 8
NUB1: NEDD8 ultimate buster 1 rVV –recombinant vaccinia virus
SCRAP: Shield-controlled recombinant antigenic protein
TUBE: Tandem Ubiquitin Binding Entities
UBL: Ubiquitin-like protein
Ub: Ubiquitin

